# A gap-junction mutation in the mouse cochlea reveals cochlear amplification is driven by outer hair cell extracellular receptor potentials

**DOI:** 10.1101/2021.08.11.455955

**Authors:** Snezana Levic, Victoria A. Lukashkina, Patricio Simões, Andrei N. Lukashkin, Ian J. Russell

## Abstract

Cochlear amplification, whereby cochlear responses to low-to-moderate sound levels are amplified and compressed to loud sounds, is attributed to outer hair cell (OHC) electromotility driven by voltage changes across the OHC basolateral membranes due to sound-induced receptor-current modulation. Cochlear operation at high acoustic frequencies is enigmatic because the OHC intracellular receptor potential (RP) is severely attenuated at these frequencies. Clues to understanding the voltage control of OHC electromotility at different frequencies was provided by measurements from CD-1 mice with an A88V mutation of the gap-junction (GJ) protein connexin 30 (Cx30), which with Cx26, form heterogeneous GJs between supporting cells in the organ of Corti (OoC) and stria vascularis. The A88V mutation results in a smaller GJ conductance which may explain why the resistance across the OoC in CD-1Cx30^A88V/A88V^ mutants is higher compared with wild-type mice. The endocochlear potential, which drives the OHC receptor current and, consequently, the OHC RPs, is smaller in CD-1Cx30^A88V/A88V^ mutants. Even so, their high-frequency hearing sensitivity equals that of wild-type mice. Preservation of high-frequency hearing correlates with similar amplitude of extracellular receptor potentials (ERPs), measured immediately adjacent to the OHCs. ERPs are generated through OHC receptor current flow across the OoC electrical resistance, which is larger in CD-1Cx30^A88V/A88V^ than in wild-type mice. Thus, smaller OHC receptor currents flowing across a larger OoC resistance in CD-1Cx30^A88V/A88V^ mice may explain why their ERP magnitudes are similar to wild-type mice. It is proposed that the ERPs, which are not subject to low-pass electrical filtering, drive high-frequency cochlear amplification.

**Significance Statement:** Cochlear amplification, whereby responses to low-to-moderate sound levels are amplified and those to loud sounds are compressed, is attributed to outer hair cell (OHC) electromotility. Electromotility is driven by voltage changes across the OHC basolateral membranes due to modulation of receptor current flow during sound-induced sensory hair bundle displacement. Mechanisms of high-frequency cochlear amplification remain to be elucidated. A mutation of the gap-junction protein connexin 30 decreases OHC intracellular receptor potentials in CD-1 mice. Instead of decreasing auditory sensitivity, the mutation rescues high-frequency hearing by causing OHC extracellular receptor potentials to be similar in amplitude to those of sensitive wild-type mice. It is proposed extracellular, not intracellular, potentials drive high-frequency OHC motility and cochlear amplification at high acoustic frequencies.

## INTRODUCTION

An intriguing challenge in auditory neuroscience is to discover how cochlear amplification, attributed to outer hair cell (OHC) electromotility (*1-5*), operates throughout the enormous frequency bandwidth of mammalian hearing (*6*). At each location, throughout its length, from low-frequency apex to high-frequency basal coil, the basilar membrane (BM), which support the sensory organ of Corti (OoC), is tonotopically tuned to a place-dependent characteristic frequency (CF). For frequencies around the CF, amplification boosts cochlear responses to low-to-moderate sound levels, while compressing them to loud sounds (*7*). OHC electromotility is driven by voltage differences generated by mechanoelectrical transducer (MET) current flow across the electrical impedance of OHC basolateral membranes, which are populated by the voltage-sensitive motor protein SLC26a5 (prestin, (*8*)). OHC motility bandwidth is potentially limited by prestin kinetics (*9*) and the OHC membrane electrical time constant (Kössl and Russell, 1992). Unlike the intracellular receptor potentials (RPs), the large extracellular receptor potentials (ERPs) recorded from the OoC are not filtered by the cell’s membrane time constant. They are, thus, are potentially sufficient to drive OHC voltage-dependent motility and provide cochlear amplification (e.g. (*10-12*)).

The CD-1 mouse strain is characterized by rapid-onset presbycusis (*13*). However, in a strain (CD-1Cx30^A88V/A88V^ mouse) that expresses the A88V/A88V mutation of Connexin 30 (Cx30), high-frequency hearing is preserved, but low-frequency hearing below ∼12 kHz is less sensitive compared with the wild-type littermates (*14-16*). This finding is surprising because the endocochlear potential (EP) in CD-1Cx30^A88V/A88V^ mice is only ∼ 70 mV compared with ∼ 112 mV found in mice with sensitive high-frequency hearing (e.g. CBA/J mice, (*15*)). EP in series with the OHC resting membrane potential (*17, 18*) provides the electromotive force that drives the OHC MET current (*19*) and EP reduction decreases cochlear sensitivity (*20*), but obviously not in CD-1Cx30^A88V/A88V^ mice.

Connexins oligomerize into hexameric structures called connexons. They form hemichannels on cell surfaces and interact with hemichannels on neighbouring cells to create gap junctions (GJs) that permit bidirectional flow of ions and signalling molecules (*21, 22*). The hemichannels of supporting cells in the OoC are formed of homotypic and heterotypic, co-localized, Cx26 and Cx30 (*21, 22*). Deletions or mutations of these connexins dominate genetically based hearing loss (*23*). Mutations of Cx30, including A88V are the basis for an autosomal dominant genetic skin disorder, Clouston syndrome (OMIM #129500) that can be associated with hearing disorders (*14*). Large GJs are formed exclusively between the OoC supporting cells, including Deiters cells (DCs) and outer pillar cells (OPCs), which encompass extracellular fluid spaces of the OoC (*24, 25*). OoC supporting cells are thought to maintain OoC homeostasis and contribute to the electrical and micromechanical environment of the sensory hair cells (*26-28*). DCs are attached to the base and apex of OHCs (*24*) and may influence OHC force production and sound-induced cochlear responses (*28-32*).

ERPs recorded adjacent to the basolateral membranes of OHCs in the OoC fluid spaces of CBA/J and CD-1Cx30^A88V/A88V^ mice are closely similar in magnitude (*15*). Our objectives were, therefore, to discover how the A88V/A88V mutation influences the electrophysiology of DCs and associated GJs, and the electrical responses and properties of the OoC, to enable ERPs be of similar magnitude in CBA/J and CD-1Cx30^A88V/A88V^ mice in which the driving voltage for the MET current is reduced. We measured the electrical properties of DCs and their GJs in situ, the intracellular RPs and ERPs from OHCs and the electrical resistance between the OoC fluid spaces and scala media *in vivo*. We investigated how these changes accounted for the sensitivity of the high frequency responses of CD-1Cx30^A88V/A88V^ mice in the face of reduced EP and if ERPs are sufficient to drive cochlear amplification at high frequencies when RPs are severely attenuated by the OHC membrane time constant.

## RESULTS

### Intracellular and extracellular cochlear electrical responses measured *in vivo*

#### Endocochlear potentials and low-frequency intracellular outer hair cell receptor potentials are larger in CBA/J than in CD-1Cx30^A88V/A88V^ mice

When gated by displacement of hair bundles in the excitatory direction, inward current flows through the MET channels at the tips of the hair cells’ stereocilia (*33*). The MET current is driven by two batteries in series due to the positive EP and the negative OHC resting membrane potential (*17, 18*), which should largely determine the magnitude of the OHC RP for a given hair bundle displacement. These RPs are predicted to be attenuated by the OHC membrane time constant (*11, 12*). This frequency limitation has been shown to be reduced, but not entirely compensated, by a tonotopic, voltage dependent potassium conductance in the OHC basolateral membranes (*34*). Intracellular RPs in response to 5 kHz tones were recorded with micropipette electrodes from presumed OHCs in the basal turns of CBA/J and CD-1Cx30^A88V/A88V^mice that had characteristic frequencies (CFs) between 52 kHz and 56 kHz. The presumed OHCs were eventually encountered after the micropipette first penetrated the BM then the supporting cells with their very negative resting membrane potentials, which are very similar in CBA/J and CD-1Cx30^A88V/A88V^ mice (CBA/J: −110.3 ± 5.4 mV, n = 15; CD-1Cx30^A88V/A88V^: −112.5 ± 4.2 mV, n = 17 (t = 1.26 P = 0.22, d.f.= 28)), and followed by the fluid-filled OoC fluid spaces with zero potential (Fig. 1A). After the micropipettes were advanced through the presumed OHCs, they entered the scala media, as indicated by a positive EP, which is significantly different (t = 47.19, P < 0.0001, d.f. = 22) between CBA/J (+112.8 ± 1.2 mV, n=12) and CD-1Cx30^A88V/A88V^ (71.3 ± 2.8 mV, n =12) mice (Fig. 1B). The presumed OHCs were identified by their large RPs and small negative resting potentials (Fig. 1B, C).

**Figure 1.**
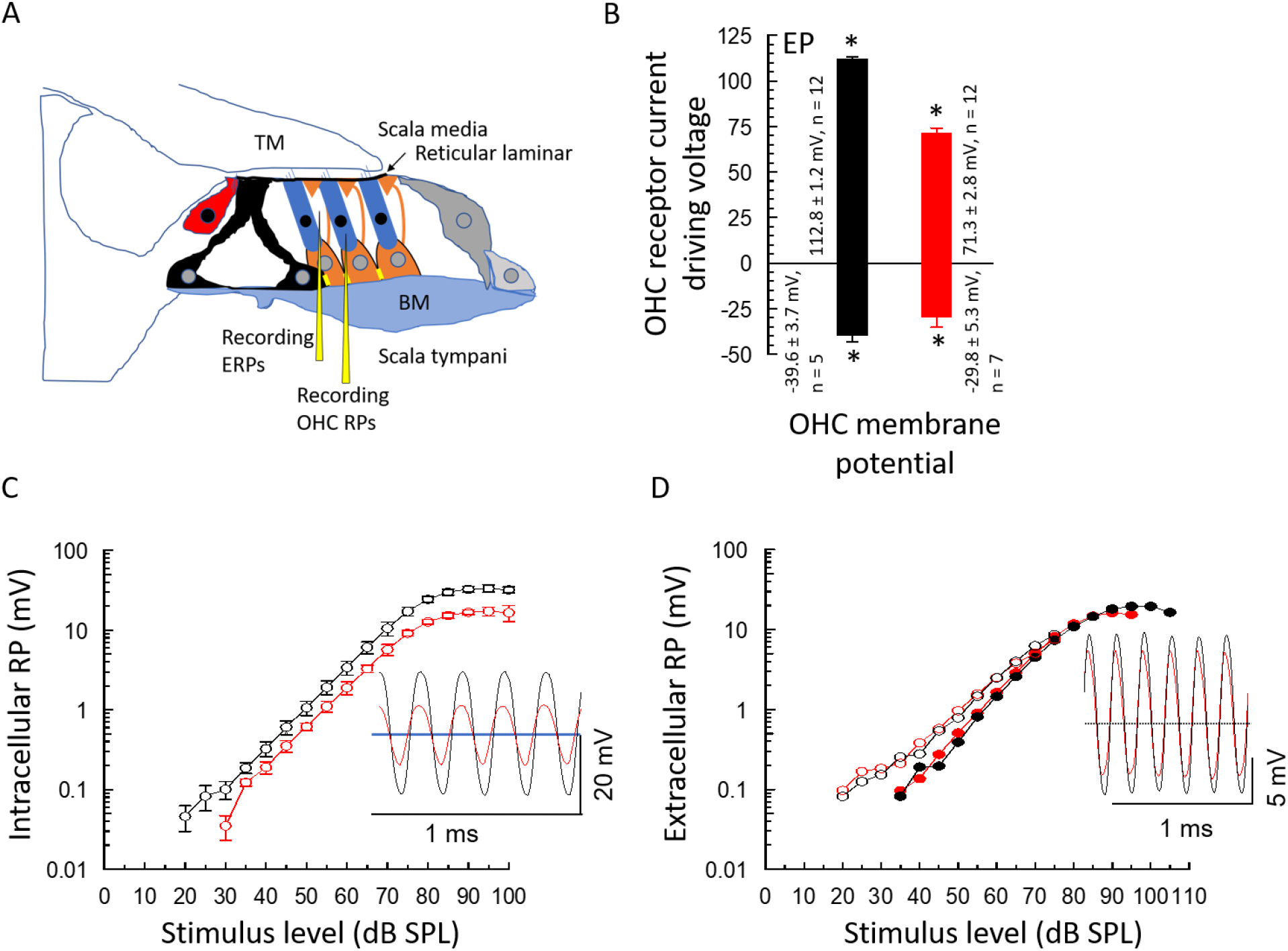
Outer hair cell intracellular receptor potential magnitudes (OHC RP) are larger in CBA/J mice (black, C) than in CD-1Cx30^A88V/A88V^mice (red, C) but extracellular receptor potential magnitudes (ERP) are similar (D). **A**. Cochlear schematic cross-section: recording arrangement for OHC RP measurements and ERP measurements from fluid spaces bounded by the OHCs (blue), DCs (orange) with phalangeal processes extending to OHC apical surface and the reticular lamina. Inner hair cells (red), pillar cells (black), basilar membrane (BM), tectorial membrane (TM). B. Mean ± SD of EP and presumed OHC resting membrane potentials for CBA/J (black) and CD-1Cx30^A88V/A88V^ (red) mice. * indicates significantly different. See text for details. **C**. Mean ± SD of the magnitude of presumed OHC intracellular RPs as functions of SPL for responses to 5 kHz tones from five CBA/J mice (black) and 7 CD-1Cx30^A88V/A88V^ mice (red). Inset: examples of intracellular RP waveforms in response to 5 kHz tones at 90 dB SPL about the resting membrane potential (blue line). Resting potentials: CBA/J, −41 mV; CD-1Cx30^A88V/A88V^, −28 mV. **D**. Magnitude of ERP from CBA/J (black) and CD-1Cx30^A88V/A88V^ (red) mice in response to stimulation with 5 kHz tones (closed symbols) and 53 kHz (open symbols) as a function of SPL. Inset: examples of ERP wave forms in response to 5 kHz, 90 dB SPL tones about the zero potential of the fluid space (dotted line). 5 kHz and 53 kHz data were taken from a CBA/J and a CD-1Cx30^A88V/A88V^ mouse with similar auditory sensitivity at 53 kHz. 53kHz data compensated for low-pass electrode characteristics (6dB octave^-1^, corner frequencies: 7.4 kHz, CBA; 9.0 kHz, CD-1Cx30^A88V/A88V^mouse).

Presumed OHC RPs recorded from sensitive CBA/J mice were large, with peak-to-peak values at 90 dB SPL of 33.3 mV (29.2 ±2.7, n = 5) and comparable in size to the largest OHC RPs recorded in guinea pigs (*35, 36*) (Fig. 1C) The largest OHC RP recorded from CD-1Cx30^A88V/A88V^mice in response to 5 kHz tones at 90 dB SPL was 18.3 mV, (17.6 ± 3.8 mV, n = 7). This is very significantly smaller than RPs recorded from presumed OHCs in CBA/J mice (t = 5.82, P = 0.0002, d.f.= 10) (Fig. 1C), although with very similar symmetry about the resting membrane potential; an indication that the operating points of the OHC transducer conductance are similar in the two mouse strains despite significant differences in EP and resting potential. The resting potentials of presumed OHCs in CBA/J mice are −39.6 ± 3.7 mV (n = 5). Those of CD-1Cx30^A88V/A88V^mice are −29.8 ± 5.3 mV (n = 7) and significantly different from those of CBA/J mice (t = 3.54, P= 0.0053, d.f. = 10, Fig. 1B). The frequency of the stimulating tones (5 kHz), which is far below that of the CF of the OHCs in the 52 kHz – 56 kHz region, was chosen because it is well within the frequency bandwidth of the micropipettes (low-pass corner frequencies 7.5 – 10.5 kHz). The magnitudes of the RPs of CD-1Cx30^A88V/A88V^ mice and CBA/J mice are close to the predicted values if the magnitude of the receptor current is proportional to the electromotive force provided by the EP in series with the OHC resting membrane potential (*17, 18*). Slight differences may indicate differences in the electrochemical environment of the OHCs between the two mouse strains.

#### Outer hair cell extracellular receptor potentials recorded in the organ of Corti of CBA/J and CD-1Cx30^A88V/A88V^mice have similar magnitudes

ERPs recorded in the fluid space immediately adjacent to the OHCs basolateral membranes are not expected to be attenuated by membrane time constants (*11, 12, 37*) and are dominated by the return flow of receptor currents from the adjacent OHCs, whose characteristics they closely share (*11*). Location of the micropipette tips in these spaces was very finely adjusted to locate them as close as possible to the basolateral membrane of an OHC without the tips being blocked through contact with cell membranes. Further advance of the micropipettes resulted either in penetration of a presumed OHC or entering the scala media. ERPs recorded from CBA/J and CD-1Cx30^A88V/A88V^ mice are not significantly different (t = 0.60, P =0.5531, df = 29) in magnitude (Fig. 1D). They are large, being maximally 17.8 mV (CBA/J:15.8 ± 2.2 mV, n = 14) and 16.5 mV (CD-1Cx30^A88V/A88V^: 15.4 ± 1.5 mV, n = 17) peak-to-peak in response to 5 kHz tones at 95 dB SPL. ERPs recorded from a CBA/J and a CD-1Cx30^A88V/A88V^mouse with similar auditory sensitivities at 53 kHz are also closely similar in magnitude and sensitivity (Fig. 1C). Thus, despite large and significant (P< 0.001) differences in the EP (Fig. 1B) and OHC resting membrane potential and RPs between CBA/J and CD-1Cx30^A88V/A88V^mice, ERPs are not significantly different.

#### The voltage difference across the outer hair cell basolateral membranes is smaller in CD-1Cx30^A88V/A88V^ than in CBA/J mice for low-frequency tones

Prestin-dependent OHC motility is understood to be controlled by the voltage difference developed across the OHC basolateral membranes (*38*). This potential was determined, in response to 5 kHz tone stimulation, for OHCs in CBA/J (Fig. 2A) and CD-1Cx30^A88V/A88V^ (Fig. 2B) mice by subtracting the ERP from the intracellular RP using the same micropipette with the assumption that the electrical time constant of the micropipette and recording system did not change during the few microns excursion of the pipette tip from fluid-filled space to the intracellular location. The ‘voltage difference’ developed across the OHC basolateral membranes are significantly larger (t = 10.40, P<0.0001, df = 10), in CBA/J mice (17.2 ± 2.6 mV, n = 5) than in CD-1Cx30^A88V/A88V^ mice (5.2 ± 1.4 mV, n = 7) peak-to-peak in response to 5 kHz tones at 95 dB SPL. The difference curves were obtained by subtracting the extracellular from the intracellular voltage responses on a point by point basis.

**Figure 2.**
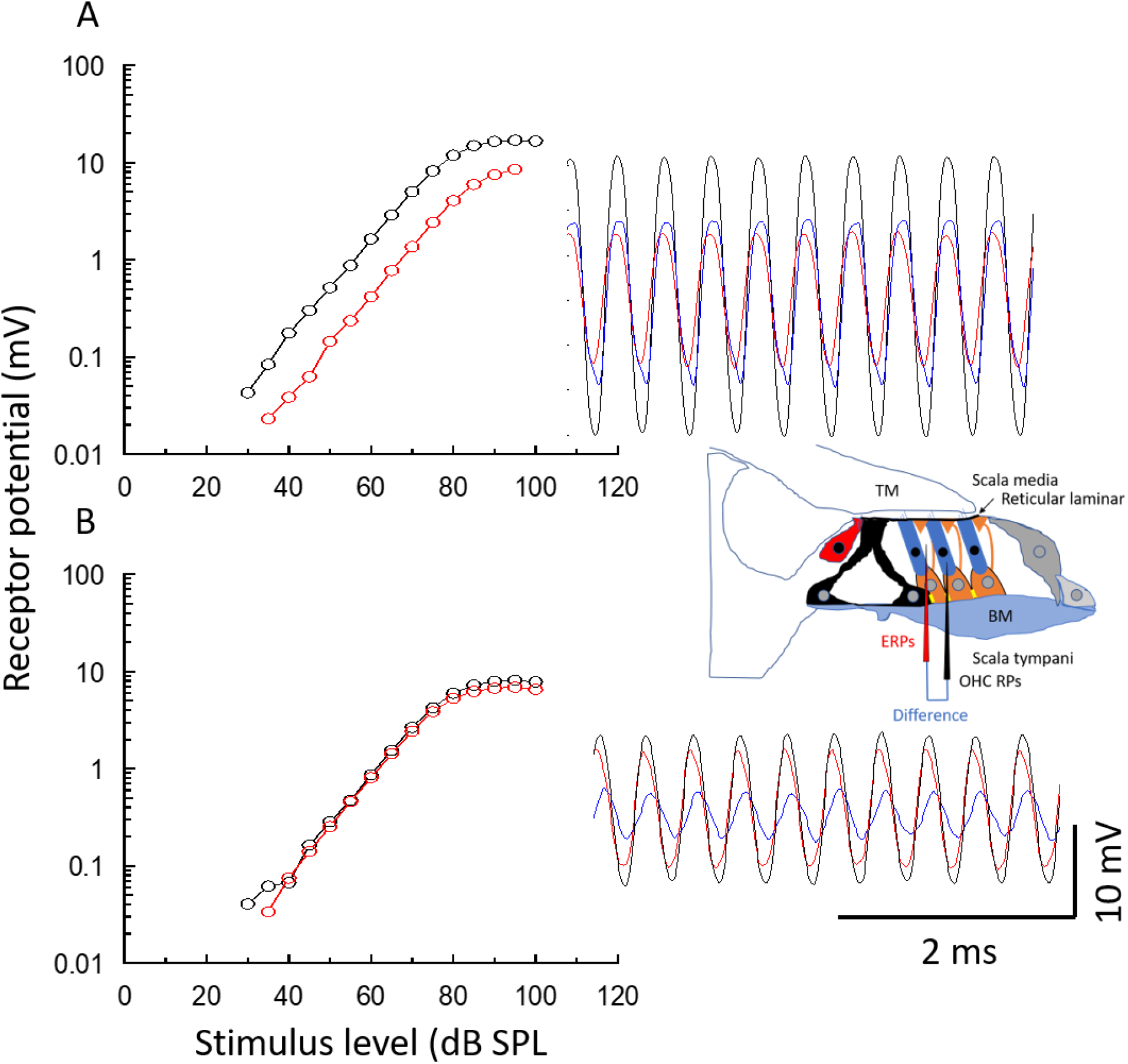
The voltage difference across OHC basolateral membranes is smaller for low-frequency tones in CD-1Cx30^A88V/A88V^ than in CBA/J mice. **A, B**, Magnitude of presumed OHC intracellular RP (open symbols) (black) and ERPs (red symbols) for CBA/J mice (A) and CD-1Cx30^A88V/A88V^ mice (B) as functions of SPL for 5 kHz tones. Insets: Schematic of recording locations and examples of waveforms recorded, in response to 5 kHz, 90 dB SPL tones, intracellularly (black), extracellularly (red), and difference (blue) for CBA/J mice (A) and CD-1Cx30^A88V/A88V^ mice (B).

#### The electrical resistance of the current path between the organ of Corti fluid spaces and the scala media is larger in CD-1Cx30^A88V/A88V^ than in CBA/J mice

How can it be that despite significantly smaller intracellular RPs in the OHCs of CD-1Cx30^A88V/A88V^ mice, the ERPs are similar in magnitude to those observed in CBA/J miceã One possibility is that the electrical properties of the OoC of CD-1Cx30^A88V/A88V^ mice are different from those of CBA/J mice. As a first step to test this hypothesis, the electrical resistance between the OoC fluid space immediately adjacent to the OHC basolateral membrane and the scala media. The micropipette tip was stepped into the OoC fluid space and most of the micropipette resistance was bridge balanced. This is necessary because the electrode resistance far exceeds that of the resistance between the scala media and OoC fluid spaces and would dominate any measurements. It was not possible to bridge balance and compensate for the microelectrode resistance completely because of its large resistance. A 100 Hz sinusoidal current from a calibrated constant current source was passed through the partially balanced resistance of the micropipette located in the OoC fluid space and immediately adjacent to presumed OHCs. The micropipette tip was then stepped a few microns through the RL into the scala media. The sinusoidal current passing across the small, unbalanced resistance of the micropipette and the electrical resistance of the current pathway between the OoC fluid space and the scala media generated a voltage that was directly proportional to the electrical resistance of this pathway. The micropipette tip was again stepped back into the OoC fluid space and sinusoidal current was again passed through the tip of the micropipette to check that the micropipette resistance remained the same. If the micropipette resistance changed, the data were discarded. The micropipette tip was finally stepped back to the scala media to discover if the penetration of the RL had caused a leak resistance (Figs. 3A, B). For all currents that provided linear voltage responses, i.e. > 5 × 10^−10^ A (see Fig. 3), we obtained the resistance by dividing the voltage responses by the injected currents. Log scales are used in Figs. 3 A, B to show that a liner relationship (see dotted lines, slopes of 1 in Figs. 3A, B) between current and voltage extended over a wide range of electrical currents. Linear scales of data presented in Figs. 3A, B are used in Figs. 3C, D to show the slope resistance of the measurements. Based on these measurements, the resistance between the OoC fluid space and the scala media, calculated as a difference of resistances measured at these two locations, was (mean ± SD) 614.2 ± 197.9 kΩ, n = 7 for CD-1Cx30^A88V/A88V^ mice and significantly smaller, 280.6 ± 37.4 kΩ, n = 8 in CBA/J mice (t = 4.70, P =0.004, d.f. = 13). This indicates the resistive pathway of the return flow of OHC receptor current, that has a major role in the determining the amplitude of the ERP (*39*), is significantly increased in CD-1Cx30^A88V/A88V^ mice.

**Figure 3.**
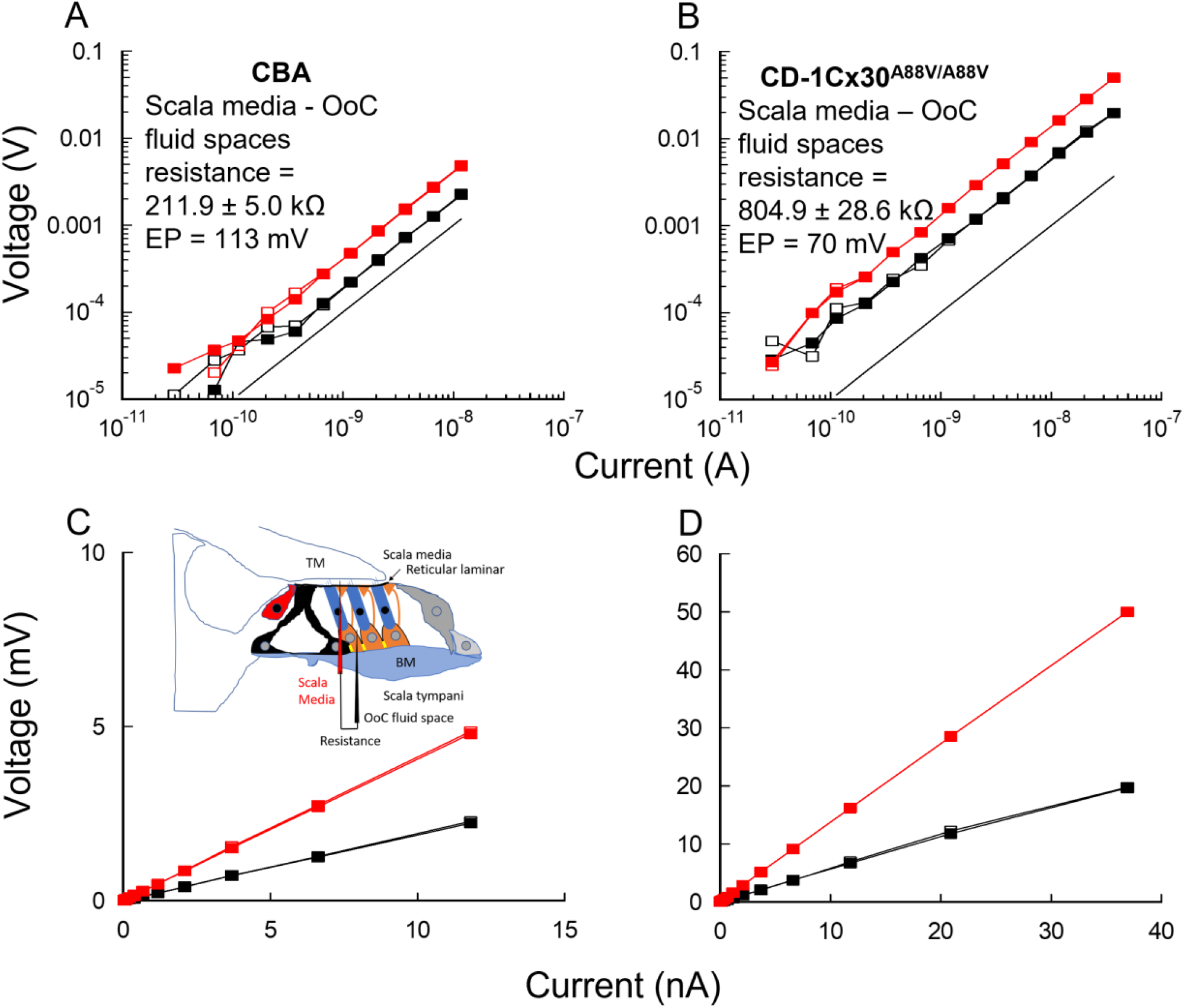
The electrical resistance of the current path between the organ of Corti (OoC) fluid space and the scala media is larger in CD-1Cx30^A88V/A88V^ than in CBA/J mice. **A, B**. Examples for CBA/J mice (A) and CD-1Cx30^A88V/A88V^ mice (B) of voltage measured (average of 20 presentations) in response to current injected through the mostly bridge-balance circuit when the micropipette tip is in the OoC fluid space next to a presumed OHC (black square) or in the scala media (red square). First measurement (open square), successive measurement when returned from scala media to OoC, or vice versa (solid square). Log-log plots to show linearity of measurement. Resistance between the scala media and the OoC fluid space calculated as the difference between resistance measurements in these two locations. C, D. Linear representations of data in A, B to show slope resistance. Inset C: Schematic to show electrode locations for the measurements in A, B. Dotted lines in A and B: slope of one.

#### Extracellular receptor potentials dominate driving voltage for outer hair cells somatic motility for frequencies above 10 kHz

In response to 5 kHz tones, the voltage generated across the basolateral membranes of presumed OHCs in the basal turn of the cochlea is larger in CBA/J mice than in CD-1Cx30^A88V/A88V^ mice (Fig. 2). However, the ERPs generated at 5 kHz and 53 kHz are similar in both CBA/J and CD-1Cx30^A88V/A88V^ mice with similar cochlear sensitivity (Fig. 1D). How does the relationship between OHC intracellular and extracellular potentials, and hence the potential difference across OHC basolateral membranes, change as a function of tone frequency? Measurements were challenging and successful recordings were made in seven CD-1Cx30^A88V/A88V^ mice, only two of which remained sensitive within 5 dB SPL of the initial thresholds throughout the 40-minute recording period. The others remained within 10 dB of the initial threshold (20 - 30 dB SPL). Measurements (mean of ten 70 dB SPL tone presentations for each frequency step within frequency range of 4-70 kHz in 2 kHz steps) were made only while the OHC membrane potential remained stable. The 70 dB SPL tone level was chosen to obtain reliable signals over a wide frequency range. This presented the problem that it was not possible to determine the true threshold (20 – 30 dB SPL) at the CF of the recording location, so the frequency that gave the largest magnitude responses was referred to as the CF. ERP (Fig. 4A) and intracellular RP (Fig. 4B) magnitudes were plotted as functions of stimulus frequency and presented as raw (open symbols) and compensated for the low-pass filter of the recording pipette measured in situ (solid symbols) (*11, 12*). The superimposed intracellular RPs and ERPs compensated for the low-pass filter of the microelectrode (Figs. 4A, B, solid symbols) are shown in Fig. 4C and for another OHC in a different cochlea in 4D. Mean data from four adjacent sets of OHC measurements from the 64 kHz region of the same cochlea are shown in Fig. 4E. In all measurements, for frequencies below ∼ 10 kHz, the magnitudes of the ERPs and intracellular RPs are similar (compare with level functions presented in Fig. 2B). For frequencies above this, the magnitude of the intracellular RP diverges from that of the ERPs with increasing frequency, presumably because of the OHC membrane time constant (*10, 11*). A magnitude notch (vertical dotted lines, Fig. 4) is most noticeable in the intracellular RPs. For measurements at BM frequency places with CFs between 58 kHz and 68 kHz (63.1 ± 3.2 kHz, n = 9), the tip of the notch is centred on frequencies between 42 kHz and 48 kHz (43.1 ± 3.2 kHz, n = 9), which is 0.552 ± 0.035 octaves below the CF of the recording site (or 0.683 CF frequency) when calculated independently for each of the nine measurement sites. The reduced size of the notch in extracellular recordings at 70 dB SPL may be due to spread of extracellular current from multiple sources and the frequency location of the notch could not be accurately determined in six of the ERP measurements. The RP magnitude increases sharply when the tone frequency is stepped above that of the notch and reaches a maximum close to the CF. The magnitude of the voltage response measured at the CF at 70 dB SPL, when compensated for the low-pass filter of the recording pipette, but not for the time constant of the presumed OHCs (resting membrane potentials −35 mV ± 4 mV, n = 4) is significantly different (t =10.10, P < 0.0001, df = 13) between the intracellular (0.64 mV ± 0.07 mV, n = 4) and extracellular (3.08 ±0.47 mV, n = 11) RPs measured from the same cochleae. The voltage difference generated across the presumed OHC basolateral membranes is indicated by the solid blue line in Fig. 4E, where it is minimal for frequencies below 10 kHz and is closely similar to the ERP for frequencies above this. Accordingly, if Prestin-dependent OHC motility is controlled by the voltage difference developed across the OHC basolateral membranes, the findings presented here indicate that OHC motility is driven by the ERP for frequencies above 10 kHz in the high-frequency region of the mouse cochlea.

**Figure 4.**
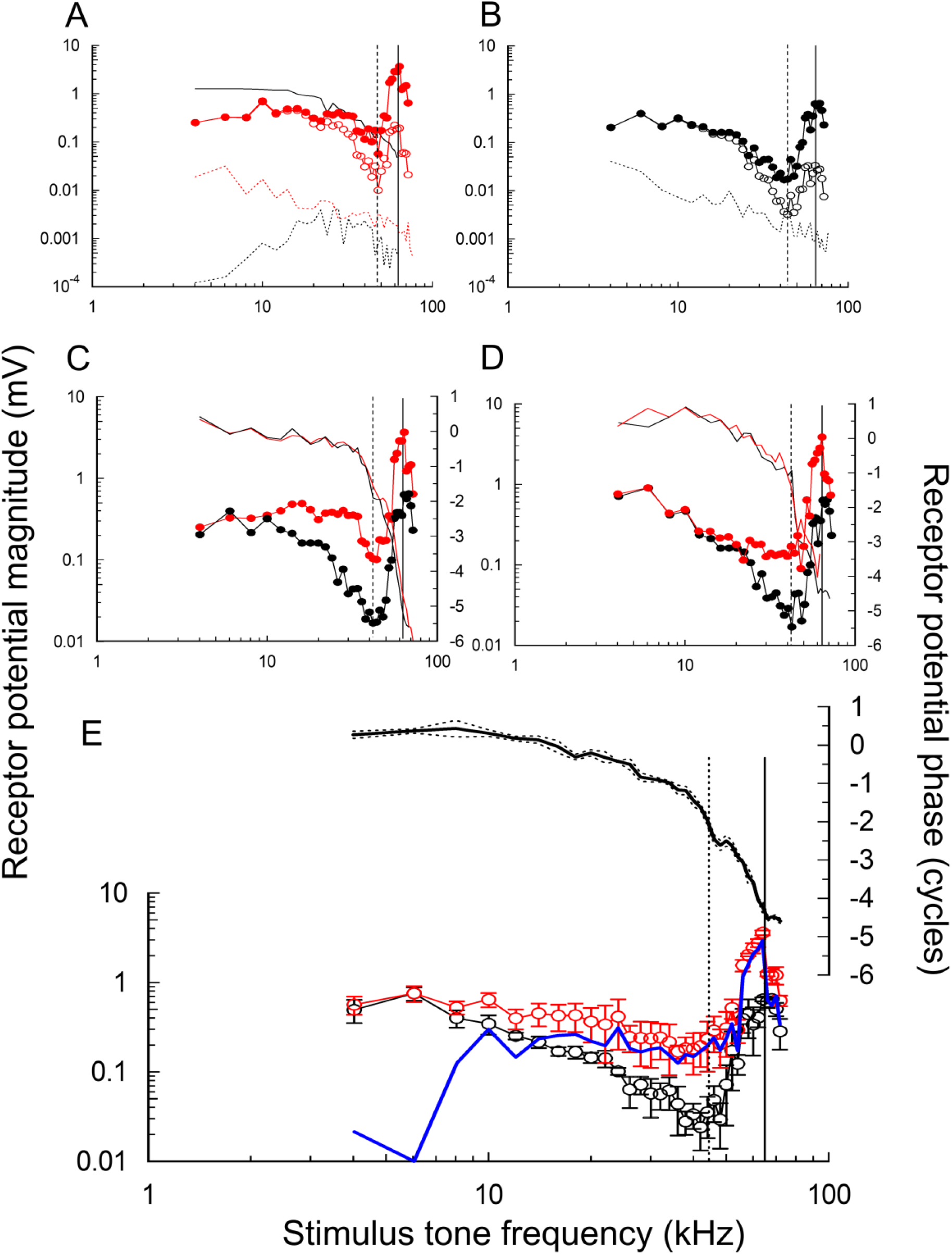
Extracellular receptor potentials dominate driving voltage for the outer hair cell somatic motility for tone frequencies above 10 kHz. A. An example of magnitude of ERP recorded in fluid space close to an OHC of a CD-1Cx30^A88V/A88V^ mouse with a CF of 62 kHz as a function of stimulus tone frequency. Open symbols: raw data. Solid symbols: data corrected for recording electrode low-pass filter characteristics (solid black line). B. Magnitude of intracellular RPs (black symbols) recorded from a presumed OHC immediately adjacent to recording in (A) (resting membrane potential – 31 mV) as a function of stimulus frequency. Open symbols: raw data. Closed symbols: data compensated for recording electrode low-pass filter characteristics. C. Superimposed, compensated data from (A) and (B) together with RP phase (solid lines) as a function of stimulus frequency. D. ERP and RPs from another OHC in a different cochlea. E. Mean ± SD of four intracellular OHC RPs and four ERP recordings made immediately adjacent to the intracellular recordings in the 64 kHz frequency region of a single cochlea of a CD-1Cx30^A88V/A88V^ mouse. Thick blue line: potential difference across presumed OHC basolateral membranes (difference between mean magnitudes of ERP and intracellular RP). Mean phase (thick black line) ± SD (dotted lines) from the four OHC RPs. Vertical solid line: CF; dotted line: notch frequency. All data from CD-1Cx30^A88V/A88V^ mice with ERP thresholds < 30 dB SPL at the CF. Phase was corrected for middle ear transfer characteristics (*40*), sound system and recording electrode. Dotted lines (A, B): recording noise floors.

Intracellular, iso-level (70 dB SPL) RP phase as a function of frequency was similar in form between measurements in different preparations with closely similar frequency locations (Figs. 4C-E). RP phase as a function of frequency is seen most clearly in the averaged data of Fig 4E. The slowly declining phase roll-off with increasing stimulus frequency becomes steeper from the onset of the notch in the amplitude iso-level frequency function. Intracellular RP phase roll-off again becomes steeper from the onset of the rising phase of the intracellular RP magnitude peak. The two frequency regions of steep phase roll-off are separated by a phase plateau for frequencies between 47 kHz and 50 kHz. This plateau represents a phase shift of 0.34 ±0.04 cycles (n=8) that is associated with the notch in the intracellular RP magnitude frequency function.

### Electrophysiological properties of Deiters cells measured *in situ*

To investigate the cellular basis for differences in the *in vivo* electrical responses that were measured in the OoC of CBA/J and CD-1Cx30^A88V/A88V^mice, potassium currents were measured in DCs from *in situ* isolated middle-turns of the cochlea of CBA/J and CD-1Cx30^A88V/A88V^ mice. CBA/J mice, with excellent high-frequency hearing, were used as the “WT control” in the *in vivo* measurements. Measurements were also made from DCs of CD-1 mice, which express rapid, early onset hearing loss, to discover if, and how, the Cx30^A88V/A88V^ mutation altered the electrophysiological properties of the DCs in the CD-1 background strain that might be associated with the preservation of high frequency hearing in the mutant mice. The ages of the mice were restricted to ∼ P20 to avoid the effects of rapid-onset-hearing loss that appear after this time in CD-1 mice.

#### CD-1Cx30^A88V/A88V^ Deiters’ cells have potassium current kinetics which are closer to CBA/J mice with good high-frequency hearing

Whole-cell currents were elicited in DCs using 500 ms depolarizing voltage steps in 10-mV increments from −90 to +50 mV, from a holding potential of −80 mV. Representative current traces are shown in Figs. 5 A, B, C. DC potassium currents were significantly different in amplitude, density, and voltage sensitive activation between CD-1, CBA/J, and CD-1Cx30^A88V/A88V^ mice (Figs. 5 D, E, F). At an activation voltage of +50 mV, the average DC potassium current amplitude of CD-1Cx30^A88V/A88V^ mice (3514 ±469 pA) was significantly larger than that of CBA/J (2829 ±416 pA) mice (t = 5.46, P< 0.0001, d.f.=48, Dunn–Sidak correction α’=0.017). However, the average DC potassium current amplitude of CD-1 mice (5430 ±724 pA) was significantly larger than either those of CBA/J, and CD-1Cx30^A88V/A88V^ mice (t = 11.11, P < 0.0001, d.f.= 48, α’=0.017). These differences were also reflected in the current density (Fig. 5E). Current density at +50 mV depolarization was significantly lower in CBA/J (270±40 pA/pF) than in CD-1Cx30^A88V/A88V^ mice (335±45 pA/pF) (t = 5.40, d.f. = 48, α’=0.017; Fig. 5D). Current density of both CD-1Cx30^A88V/A88V^ and CBA/J was significantly lower than in CD-1 mice (517±68 pA/pF; t = 11.16, P < 0.0001, d.f.= 48, α’=0.017; Fig. 5E). Differences in current density were not due to differences in cell size because cell capacitance was similar in CD-1 CD-1Cx30^A88V/A88V^, and CBA/J being 11.3 ± 0.4 pF, 11.6 ± 0.5 and 11.4±0.6, respectively (n= 149). The current traces in Figs. 5 A, B, C indicate that the potassium conductance shows little inactivation in the DCs of CD-1 mice and in contrast with the increasing levels of inactivation observed in the DCs of CD-1Cx30^A88V/A88V^ and CBA/J mice. Voltage dependence of activation was shifted to more negative potentials for potassium currents in CD-1Cx30^A88V/A88V^ and CBA/J compared to that of CD-1 DC’s (Fig. 5F). The half activation voltages (V_1/2_) occurred at significantly (t = 6.81, P < 0.0001, d.f.= 48, α’=0.017) lower potentials in CD-1Cx30^A88V/A88V^ (V_1/2_ =12.4 ± 2.8 mV) DCs than in CBA/J (V_1/2_ =16.5 ± 1.1 mV) DCs. These potentials in the DCs of CD-1Cx30^A88V/A88V^ were significantly smaller (t = 22.36, P < 0.0001, d.f.= 48, α’=0.017)) from those measured in the DCs of CD-1mice (V_1/2_ = 27.9 ±2.3 mV).

**Figure 5.**
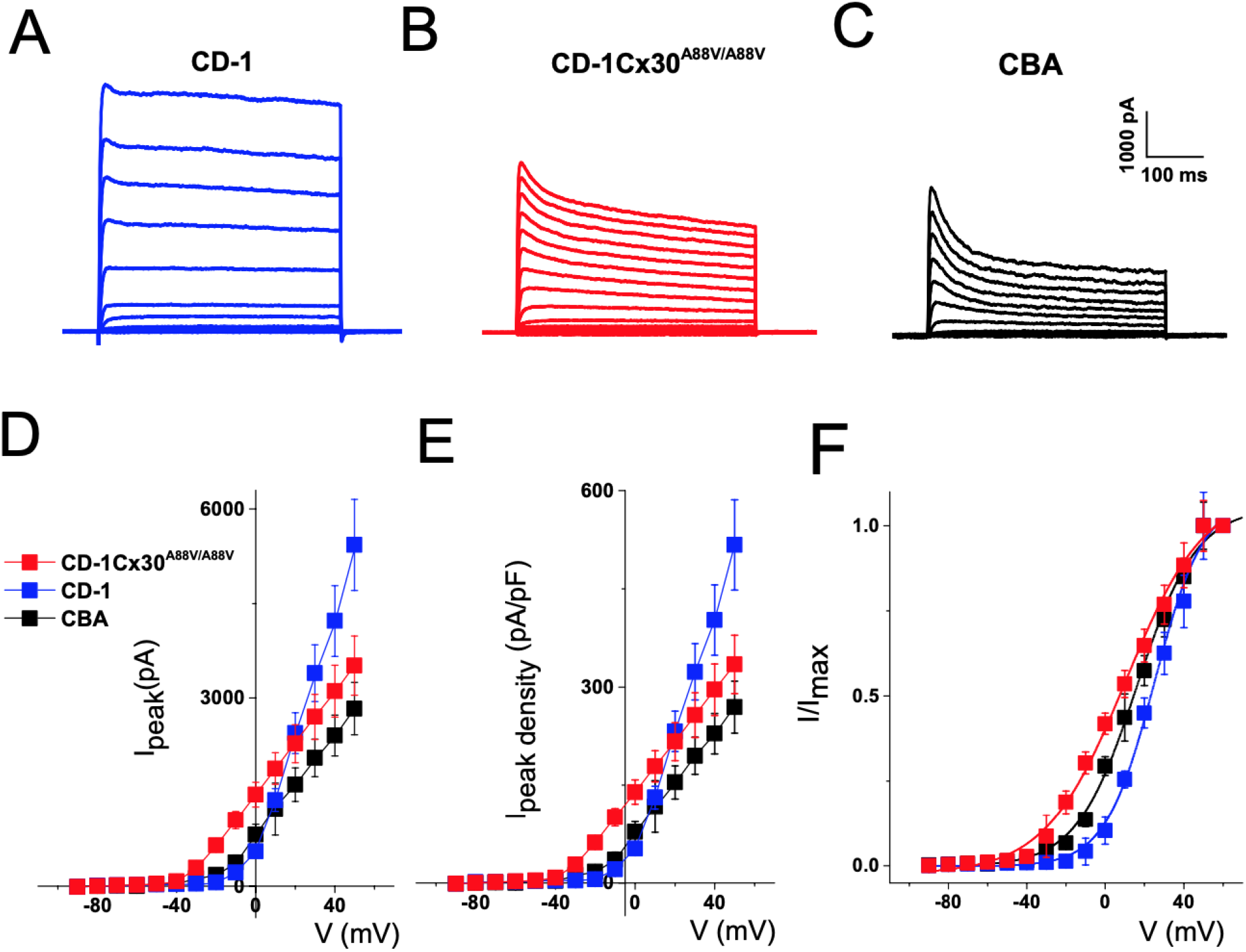
Potassium current amplitude, density and voltage sensitive activation are altered in CBA/J and CD-1Cx30^A88V/A88V^compared to CD-1 Deiters’ cells (DCs). Representative examples of current traces recorded from CD-1 (A), CD-1Cx30^A88V/A88V^ (B) and CBA/J (C) DCs. The current-voltage (D) and density-voltage (E) relationships are significantly different between CBA/J, CD-1 and CD-1Cx30^A88V/A88V^ DCs. (F) The voltage dependence of activation was assessed for peak potassium currents elicited in CD-1 and CD-1Cx30^A88V/A88V^ DCs. The Boltzman fits are plotted as solid lines. The half-activation voltages (V1/2) were more positive for CD-1 cells (27.9 ± 2.3 mV) than for CD-1Cx30^A88V/A88V^ cells (12.4 ± 2.8 mV). CBA/J mice were in between (half-activation voltage (V1/2) was 16.5 4 ± 1.1) The maximum slope factor (k) was 13.6 ± 1.2 for CD-1, 21.3 ± 2.0 for CD-1Cx30^A88V/A88V^ and 15.3 ± 0.9 for CBA/J DCs. Each point (mean ± SD.) is based on measurements for n = 25 cells.

Together, our results show distinct differences in DC potassium conductances between CD-1Cx30^A88V/A88V^ and CBA mice that are both significantly different from those in the DCs of CD-1 mice, with early onset hearing loss (Fig. 5).

#### Heterogeneous transjunctional voltage coupling between Deiters cells was observed in CD-1 and CBA/J but not in CD-1Cx30^A88V/A88V^mice

Previous studies demonstrated that, in addition to homeotypic GJ conductance (HomC), most GJ conductance in the OoC and cochlear partition have asymmetric voltage gating, which has been taken to indicate heterogeneous coupling (HetC), perhaps resulting from heterotypic channels or possibly heteromeric configurations (*41, 42*). Pairs of adjacent DC pairs were initially clamped to −40 mV. Transjunctional voltage (V_j_) was produced by 2-second voltage steps applied to cell 1 from −110mV to +60 mV (10 mV increments), and cell 2 continuously held at −40 mV. The change in V_j_=V_2_-V_1_ was thus from −100 to +70 mV. Typical examples of transjunctional currents (I_j_) recorded during 1s pulse stimulation from pairs of CD-1 and CD-1Cx30^A88V/A88V^ DCs are shown in Fig. 6A and for CBA/J in Fig. S1.

**Figure 6.**
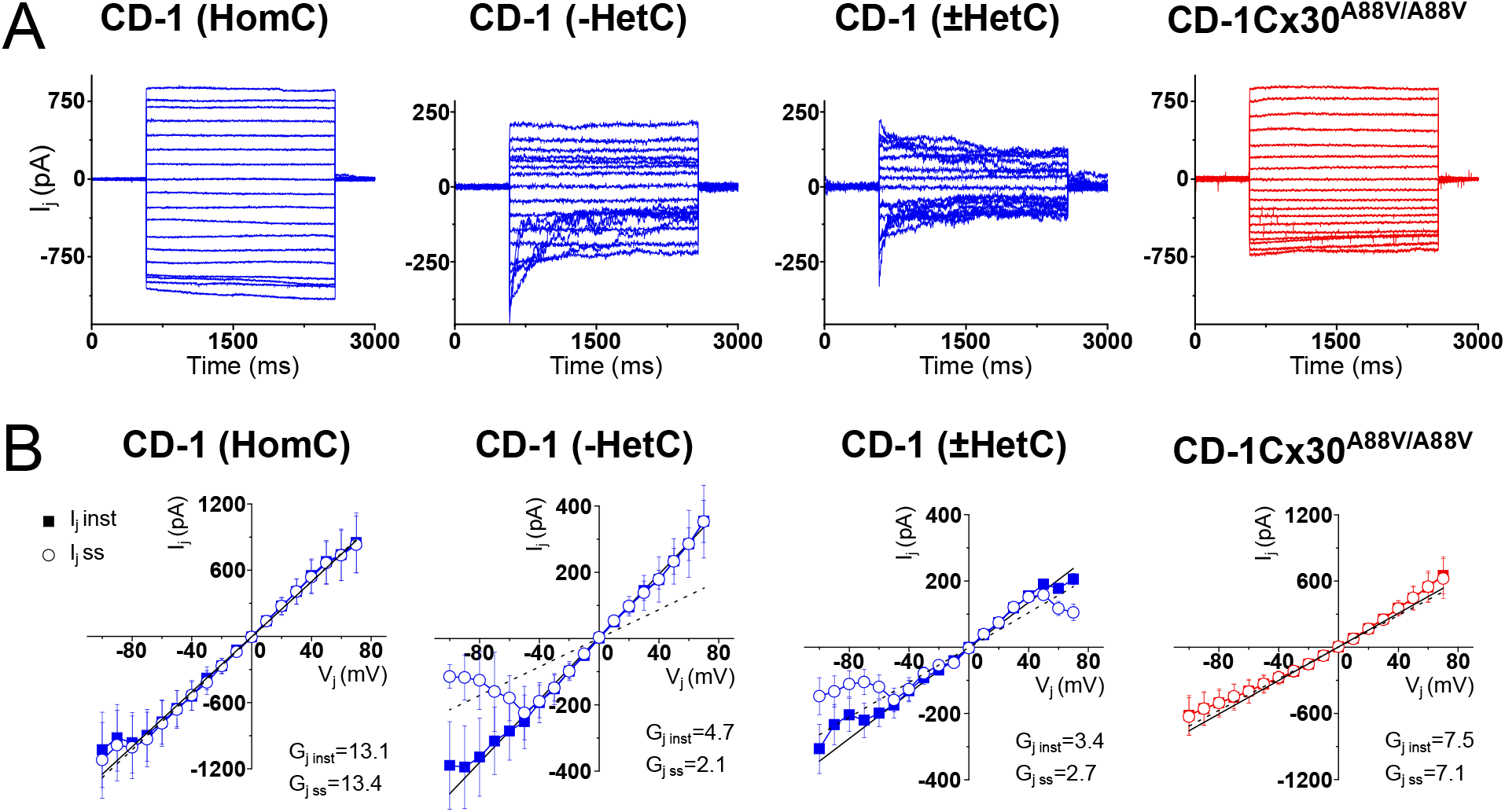
Heterogeneous transjunctional voltage coupling was observed in CD-1 and CBA/J (see S1) but not in CD-1Cx30^A88V/A88V^ Deiters cells (DCs). (A) Typical examples of transjunctional currents (I_j_) recorded during 2-sec pulse stimulation from pairs of CD-1 and CD-1Cx30^A88V/A88V^ DCs. Both cells were initial held at −40 mV and step voltage was applied to one cell and the transjunctional currents were measured in the non-stepped adjacent cell. CD-1 but not CD-1Cx30^A88V/A88V^ DCs showed a heterogenous transjunctional voltage (V_j_) and time dependence, defined by three different categories of response (HomC, -HetC, ±HetC). (B) Summary plots of I_j_ in response V_j_ stimulation obtained from the DC pairs, as classified above. Instantaneous (solid squares) and steady-state (open circles) I_j_ were measured at onset (5 ms) and end (∼2 sec), respectively (showing mean ± SD, n = 6 for each panel). Instantaneous and steady-state gap junctional conductances (G_j_) values correspond to the slope of the linear regression of the represented I_j_-V_j_ relationships.

Transjunctional currents (I_j_) in the DCs of CD-1Cx30^A88V/A88V^ mice are homogenous with little time-dependence (Fig. 6A). In contrast, I_j_ in CD-1 and CBA/J DCs displayed heterogenous voltage and time dependence. We were able to separate the DC responses of CD-1 mice into 3 categories based on I_j_, voltage and time dependence: CD-1(HomC), where the I_j_ displayed no or little voltage- and time-dependent inactivation; CD-1(-HetC) shows I_j_ with pronounced voltage- and time-dependent inactivation when the V_j_ is lower than −50 mV; and CD-1 (±HetC) showed I_j_ with pronounced voltage- and time-dependent inactivation observed for V_j_ more negative than −50 mV and more positive than 50 mV (Fig. 6A). These two last categories also showed lower I_j_ magnitudes than CD-1 and CBA/J (HomC). Figure 6B shows the summary plots of I_j_ in response to V_j_ stimulation obtained from DC pairs using the above classification. The peak instantaneous (I_j.Inst_) and steady-state (I_j.SS_) transjunctional currents were measured at the onset (∼5 ms) and the end of each voltage step (∼2 sec), respectively. In all cell types, the I_j.Inst_ relative to the V_j_ was approximately linear; I_j.SS_ showed, however, heterogenous deviation from linearity in CD-1 and CBA/J cells for potentials more negative than −50 mV (in CD-1 and CBA/J (-HetC)) and more positive than 50 mV (in CD-1 and CBA/J (±HetC)).

The comparison of the instantaneous conductances of DCs across different strains (Fig. 7) shows that the conductance of non-inactivating I_j_ in CD-1Cx30^A88V/A88V^ is 40-50% of that of CD-1(HomC). Measured at +70 mV, conductance was (mean ± SD, n = 6) 8.9 ± 0.7 nS for CD-1Cx30^A88V/A88V^ and 12.5 ± 1.2 nS for CD-1(HomC). CD-1Cx30^A88V/A88V^(HomC) conductance is, however, 110-160% higher than conductance for inactivating I_j_ (CD-1(-HetC) and (±HetC)). Measured at +70 mV, conductance was (mean ± SD, n = 6) 5.1 ± 0.6 nS for CD-1(-HetC) and 2.9 ± 0.3 nS CD-1(±HetC). At the DC resting membrane potential (∼110 mV), CBA/J(HomC) conductance, which is observed in about 75% of the DC pairs, is twice as large as CD-1Cx30^A88V/A88V^ GJ conductance.

**Figure 7.**
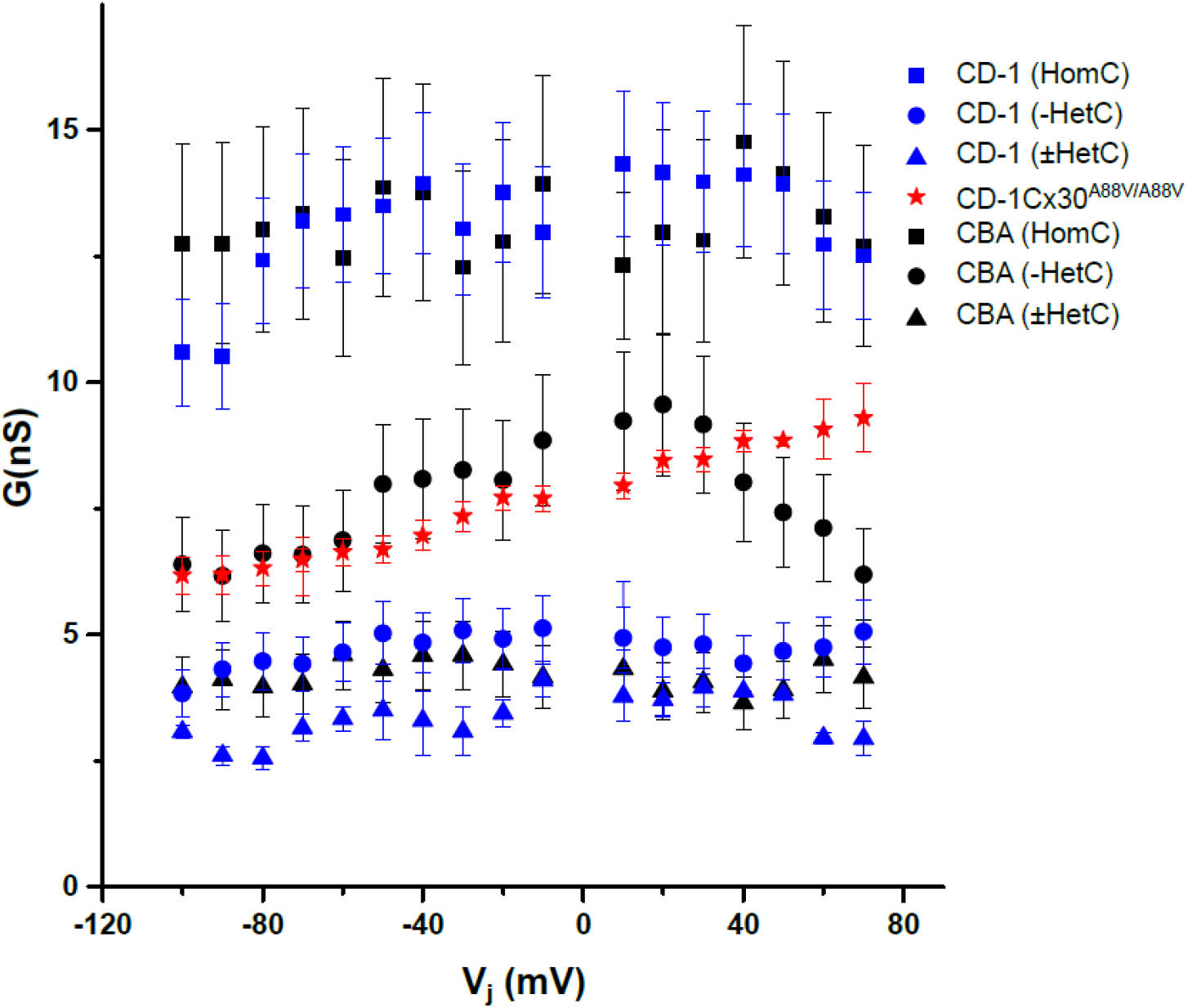
Summary comparison of Deiters cells’ conductance values in CD-1, CBA/J and CD-1Cx30^A88V/A88V^ cells as determined from I_j_ vs V_j_ instantaneous plots. Voltage independent couplings (CBA/J and CD-1 (HomC)) have the highest conductance, which is about 3 times higher than the voltage dependent conductances (CBA and CD-1 (-HetC) and (±HetC)). Although, CBA/J DCs had higher conductances than CD-1 DCs with similar responses. Mean ± SD is shown, n = 6 for all data points.

#### Vj-dependent gating in pairs of Deiters cells produces symmetrical responses in CD-1Cx30^A88v/A88V^ but heterogeneous responses in CD-1 and CBA/J strains

The transjunctional currents of double patched DC pairs were investigated by first stimulating one cell in a pair with voltage steps and recording the I_j_ in the second cell, and then stimulating the second, while recording in the first cell of the pair (Fig. 8, with additional examples in Fig. S2) The response of CD-1Cx30^A88V/A88V^ DC pairs to transjunctional voltage reciprocal stimulation typically produced homogenous and almost symmetrical I_j,_ showing minimal time dependence. The response of CD-1 and CBA/J DC pairs to transjunctional voltage reciprocal stimulation was heterogenous, with some pairs producing strong, symmetrical I_j_, whereas other pairs produced asymmetrical responses with V_j_-dependent rectifying currents. These results suggest that the connexin channel composition is heterogeneous between CD-1 and CBA/J DC pairs, whereas CD-1Cx30^A88V/A88V^ DC pairs form homogeneous GJs.

**Figure 8.**
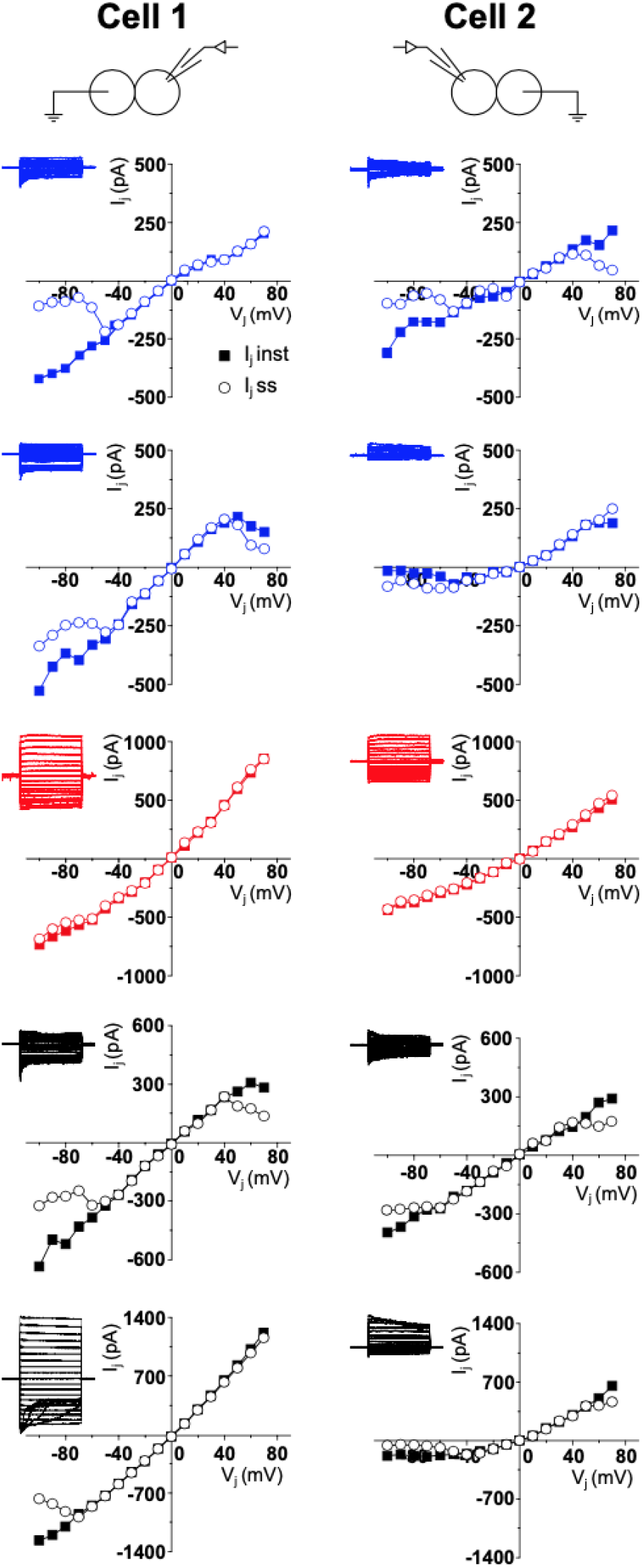
Vj-dependent gating in the same cell pair produces heterogeneous responses in CD-1 (blue) and CBA/J (black) Deiters cells (DCs) and symmetrical responses in CD-1Cx30^A88V/A88V^ (red) cells. Examples of responses elicited in different coupled DC pairs. (Cell 1) Plots of I_j_ in response to V_j_ stimulation elicited in cell 1 in response to voltage step stimulation of cell 2, and (Cell 2) response of cell 2 to voltage step stimulation in cell 1. Instantaneous (I_j.Inst_, solid squares) and steady-state (I_j.SS_, open circles) were measured at the onset (∼5 ms) and the end of each voltage step (∼2 sec), respectively.

## DISCUSSION

### Significance of Deiters’ cell electrophysiology, gap junctions and cochlear partition resistance for the magnitude of extracellular outer hair cell receptor potentials in Cx30^A88V/A88V^ and CBA/J mice

An objective of this paper is to discover how, despite having a strongly reduced EP (Fig. 1B) and, hence, a reduction in the electromotive force that drives the OHC MET current (*17, 18*), CD-1Cx30^A88V/A88V^ mice, like CBA/J mice, but unlike their wild-type CD-1 littermates, have sensitive high-frequency hearing (*14-16*). Sensitive high frequency hearing appeared to be more remarkable when it was discovered that, in response to 5 kHz tone stimulation, OHC intracellular RPs are smaller in CD-1Cx30^A88V/A88V^ than in CBA/J mice, presumably due to reduced EP. Remarkably, magnitudes and sound-pressure level-dependency of ERP are similar in Cx30^A88V/A88V^ and CBA/J mice, despite large differences in the magnitudes of the EP and OHC RPs. Measurements of the electrical resistance (R_FS_, Fig. 9) across the RL between the scala media and OoC fluid space, which in Cx30^A88V/A88V^ is over double that in CBA/J mice, provide a possible explanation for this difference. Due to increased resistance between the scala media and OoC fluid space, a given current flow across this resistance should generate a larger voltage drop in Cx30^A88V/A88V^ than in CBA/J mice. From the electromotive force driving the MET current (∼100 mV in Cx30^A88V/A88V^ mice and ∼150 mV in CBA/J mice), and the resistance between the scala media and OoC fluid space (Cx30^A88V/A88V^ mice = 624 kΩ; CBA/J mice = 281 kΩ), smaller extracellular receptor currents should be needed in Cx30^A88V/A88V^ mice to generate ERPs of closely similar size to CBA/J ERP. For Cx30^A88V/A88V^ mice, and from Ohms law, the receptor current flowing across a 624 kΩ resistance to generate an ERP of 16.5 mV is 26.8 nA. A receptor current of 63.3 nA is required to generate ERPs of 17.8 mV across a 281 kΩ resistance in CBA/J mice. Thus, in Cx30^A88V/A88V^ mice a smaller receptor current flowing across a larger resistance in the OoC produces an ERP in the OoC fluid space similar in magnitude to that recorded in the same location of CBA/J mice (Fig. 9).

**Figure 9.**
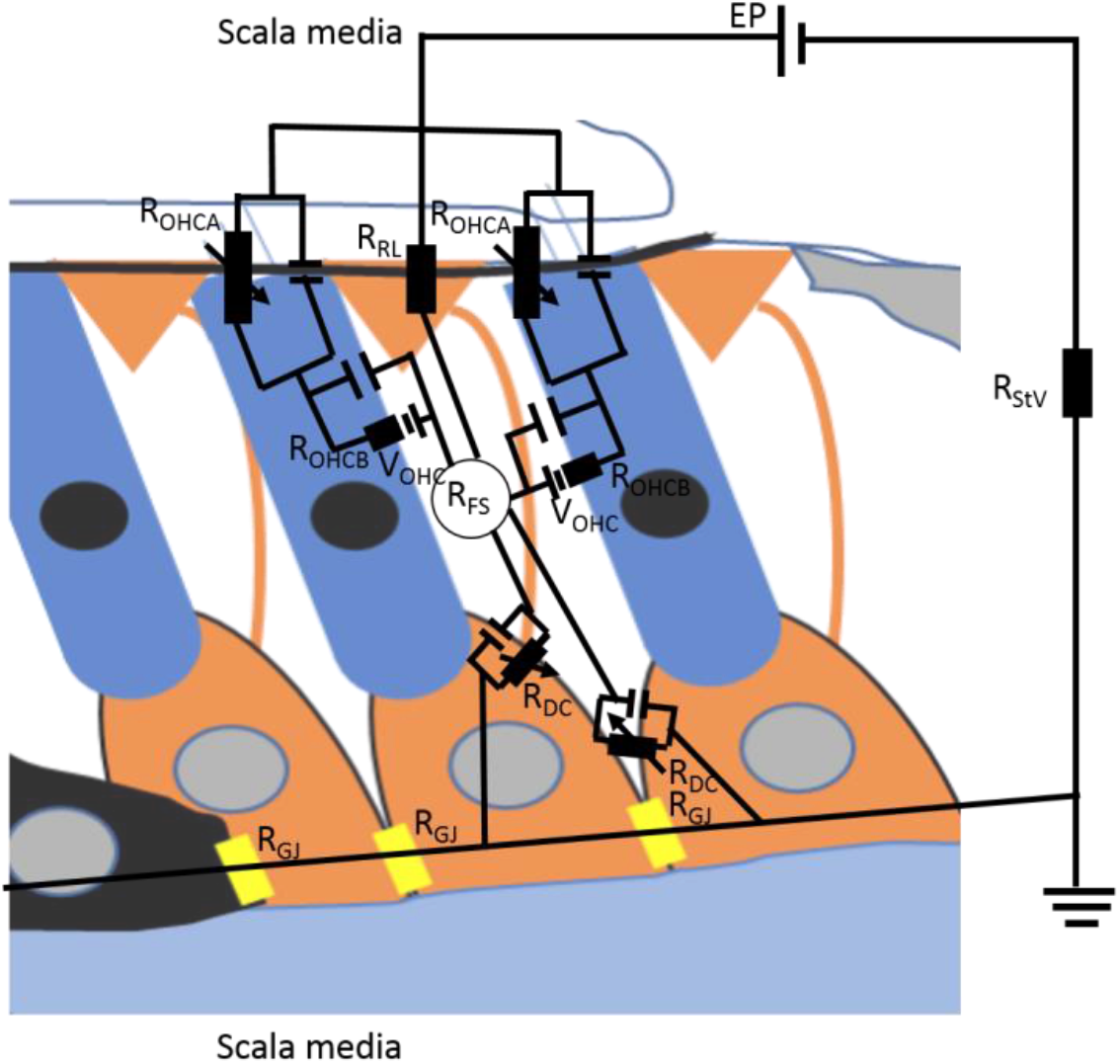
Influence of the Cx30 A88V mutation on the resistance network contributing to the electrical impedance (R_FS_) of the OHC extracellular fluid space. It is assumed that the A88C mutation of Cx30 has no direct impact on the OHC membrane conductance, indicated by R_OHCA_ for the MET conductance and R_OHCB_, for the conductance of the basolateral membrane. It is also proposed that the resistance of the reticular lamina (R_RL_) is unaffected by the mutation. The mutation reduces the overall conductance of the GJs that electrically couple the supporting cells of the OoC and stria vascularis, thereby increasing their effective resistance (R_GJ_, R_StV_) and that of the overall return pathway of the MET current through the OoC and stria vascularis. The batteries responsible for driving the MET current, the EP in series with the OHC membrane potential (V_OHC_), are both reduced by the Cx30 mutation together with the MET current. In summary, due to the Cx30 A988V mutation, resistances associated with the supporting cells in this network are increased, the driving voltages for the MET current are decreased and the MET current is decreased, but a smaller MET return current flowing across a larger OHC extracellular fluid spaces resistance (R_FS)_ results in extracellular RPs (ERP) in Cx30^A88V/A88V^ and CBA/J mice (without the Cx30 A988V mutation) being of similar magnitude.

We propose the resistance increase is due to altered current pathways within the OoC contributed partially or entirely as a result of altered electrophysiological properties of DCs and interconnecting GJs due to the Cx30^A88V/A88V^ mutation. The GJ conductance between DCs expressing the Cx30^A88V/A88V^ mutation is less than a half of the dominating homogeneous GJ conductance between DCs in CBA/J mice. In support of this proposal, the size of supporting cell Cx30 GJs in Cx30^A88V/A88V^ mice and overall expression of Cx30 GJs is reduced in the cochlea (*16*). Expression of A88V Cx30 in *in vitro* cultured cells and cell pairs causes a reduction in the conductance of GJs and their voltage gating properties (*43, 44*). Our results suggest that, heterogeneous GJ coupling was absent (*41, 42*) and GJ conductance is determined only by homogeneous GJ coupling in DCs from Cx30^A88V/A88V^mice. Positive EP is generated in the cochlear lateral wall by marginal cells and the intermediate cells coupled with the basal cells in the stria vascularis and neighbouring fibrocytes in the spiral ligament by GJs (*45*). Loss of heteromeric GJs in these epithelial cells is suggested to cause reduced EP (*46*), as also observed in Cx30^A88V/A88V^ mice. Moreover, inactivating potassium currents, that are prevalent in DCs of CBA/J and CD-1Cx30^A88V/A88V^ mice with excellent hearing are absent from the DCs of CD-1 mice with early onset hearing loss. Their significance for early onset hearing loss and its treatment are targets of our further investigation.

### The significance of the relative magnitudes of the intracellular and extracellular outer hair cell receptor potentials for cochlear amplification

The cellular basis for cochlear amplification, OHC somatic electromotility (*3-5*), is controlled cycle-by-cycle by the potential difference developed across the OHC basolateral membrane (*38*), which occurs when the cochlea is excited by sound (reviewed by (*4, 19*)). From *in vivo* measurements reported here, this potential is the frequency-dependent difference between the OHC intracellular RP and the ERP. In response to a tone of 5 kHz, which is far below the CFs of the OHCs at the recording site in the basal turn of the cochlea (52 kHz – 66 kHz), the intracellular OHC RP is larger than the ERP and dominates the transmembrane potential difference across the OHC basolateral membrane in CBA/J mice with excellent hearing. OHC intracellular RPs are considerably smaller in Cx30^A88V/A88V^ than in CBA/J mice. Consequently, the transmembrane potential difference is also smaller. If this situation pertains to OHCs in the 5 kHz location of the cochlea apical turn, it could account for the finding that for tone frequencies below 12 kHz, the auditory sensitivity of Cx30^A88V/A88V^ mice is less than that of their wild-type CD-1 littermates and sensitive CBA/J mice (*14-16*). With increasing frequency above ∼ 10 kHz, the OHC transmembrane voltage which controls the electromotility of OHCs, and hence cochlear amplification, is dominated by the ERP. This might be anticipated because it has been shown that intracellular, but not extracellular, RPs are subject to low-pass filtering of the OHC membrane time constant, which leads to attenuation of the intracellular RP by 6dB/octave above the membrane’s corner frequency (*10, 11, 37*). *In vitro* measurements demonstrate that the OHC membrane time constant is CF dependent (*34*). Even so, this dependence does not appear to fully compensate for the attenuation of OHC voltage responses to high-frequency tones. Through inspection and extrapolation of their figure 7, the intracellular RP at 53 kHz, is predicted to be reduced by more than an order of magnitude compared to that at 5 kHz for the same peak-peak receptor currents. In fact, the OHC intracellular RPs reported here are slightly larger than might be anticipated from extrapolating electrical measurements of isolated OHCs (*34*). The larger RPs and the relatively depolarised OHC resting membrane potentials, which are significantly more positive in presumed OHCs of CD-1Cx30^A88V/A88V^ than in CBA/J mice, could be due to the introduction of a leak conductance when inserting the intracellular electrode, or to other factors including the tonotopic increase in both the OHC MET conductance and the voltage dependent potassium conductance (*34, 47*). OHC resting membrane potentials are determined by the balance between the depolarizing MET current and hyperpolarizing KCNQ4-based potassium conductance of the basolateral membranes (*34, 48*). Increased activation of the voltage-dependent potassium conductance and increased MET conductance should together cause a reduction in the low-pass filtering of the OHC membrane time constant and increase in the intracellular RP, but not sufficiently to compete with the ERPs for the control of the OHC transmembrane potential and cochlear amplification for tone frequencies above 10 kHz. Similar findings of the frequency independence of extracellular potentials recorded in the OoC are reported for the guinea pig cochlea, at least for frequencies below 24 kHz (*10, 11*). The findings, reported here, support the hypothesis that intracellular receptor potentials dominate the voltage control of OHC motility for frequencies below about 12 kHz in the mouse cochlea. For frequencies above this cochlear amplification is controlled by the ERP.

### The notch in the receptor potential iso level response, the tectorial membrane, and the timing of cochlear amplification

The timing of the relative movement of structures within the cochlear partition is critical for cochlear amplification. To be effective in controlling cochlear amplification, OHC voltage-dependent forces, controlled by the ERPs and fed back to the cochlear partition, should be in phase with the BM velocity (*49*). At frequencies and levels where they are most effective, forces generated by the OHCs can enhance the BM’s motion 10000-fold at threshold (*7*). Timing of the OHC excitation is determined by the relative movement between the RL, from which stereocilia, the mechanosensitive organelles of the OHCs protrude, and the TM in which the stereocilia are imbedded. There is substantial theoretical, indirect and direct evidence to support the hypothesis that the TM coupled to the OoC acts as a resonant structure which determines the timing of the delivery of OHC electromotility that is critical for cochlear amplification (*50-56*). The amplitude notch and associated phase jump in the ERPs and intracellular RPs recorded from OHCs reported here, which occurs about a half octave below the CF, supports earlier (*57*) and current findings (*58*) about the multimodal vibrations involving the TM. The frequency of the amplitude notch indicates the resonance of the TM when the radial stiffness and mass impedance of the TM are equal and cancel each other, hence, minimizing the OHC excitation and producing a local minimum of the OHC RPs and ERPs. At this frequency, as reported here, intracellular RPs show an advance in phase of ∼ 0.3 cycles, which would introduce the correct phase relationship between mechanical and electrical responses of the cochlea to enable cochlear amplification (*49*). Indeed, at this frequency and above, the amplitudes and phases of OHC and BM responses show the level-dependent properties associated with cochlear amplification (*11, 59*). The notch in the iso-level, frequency, OHC RP function is more sharply tuned than when measured extracellularly, when it becomes increasingly broader and less distinct with increasing levels (*58*). We attribute this difference to the level-dependent increasing numbers of generators (OHCs) contributing to the extracellular signal (*60*) that smears the phase data rather than to damping by fluid in the subtectorial space, as has been recently suggested and modelled based on cochlear microphonic measurements (*58*).

We demonstrated that the ERP dominates the driving voltage for OHC somatic motility at the CF in the basal, high-frequency region of the mouse cochlea (Fig. 4). Intracellular RP, however, dominates the driving voltage when the same cochlear region is stimulated with frequencies below the cut-off frequency of the OHC membrane (Fig. 2A). If the last observation is true for the low-frequency cochlear apex, and the intracellular RP dominate the driving voltage for the OHC somatic motility in this region, it implies very different phase relationships (probably close to counter phasic) between the OHC excitation and force generation and amplification mode at the base and apex of the mouse cochlea (*61*). Attempts to relate OHC intracellular and extracellular measurements to the movements of the BM have been made (*12, 59*), but the measurements reported here, and others made previously in the guinea pig cochlea (*12, 36*) emphasise the importance of comparing intracellular OHC RPs and ERPs with measurements from the other OoC structures and from the TM (*28, 62-66*) to discover their relative contribution to the passive mechanics, amplification and frequency tuning in the cochlea.

## MATERIALS AND METHODS

### Animal preparation

#### In vivo measurements

Mice at 3-5 weeks of age were anesthetized with urethane (ethyl carbamate; 2 mg/g body weight i.p.) for surgical procedures at the University of Brighton. Mice were tracheotomized, and their core temperature was maintained at 38 °C. A caudal opening was made in the ventro-lateral aspect of the right bulla to reveal the round window. The right parietal and intarietal bones were firmly secured to the head holder using a stainless-steel rod and dental acrylic cement.

#### In vitro measurements

Mice from the same colony as those used for the *in vivo* measurements were culled using schedule 1 method.

All experiments complied with the guidelines from the Home Office under the animals (Scientific Procedures) Act of 1986 and were approved by the University of Brighton Ethical Review Committee.

### In vivo measurements

#### Electrophysiology

An Ag/AgCl ground electrode was inserted into the muscles of the neck. Intracellular electrodes (40–60 MΩ, 3 M KCl filled) were pulled from 1 mm O.D., 0.7 mm I.D. quartz glass tubing on a Sutter P-2000 micropipette puller. Signals were amplified and conditioned using laboratory-built pre-amplifiers and conditioning amplifiers that were carefully isolated and screened to avoid electromagnetic contamination from the sound system. The resistance and capacitance of the pipettes acts as a low-pass filter with a corner frequency that shifts to lower frequencies with the depth of penetration into the OoC. The frequency responses of the electrodes were, therefore, calibrated in situ according to a technique first described by (*10, 12, 67*). The filter behaved as a single-pole low-pass filter, with 3 dB cut-off frequencies ranging from 4.5 to 9.5 kHz without capacitance compensation. Electrodes were advanced using a piezo-activated micropositioner (Marzhause GMBH). The pipette tip was inserted through the round window membrane and into the BM under visual control to a location close to the feet of the OPCs. The first cells to be encountered had resting potentials of ∼ −100 mV, could be held in stable condition for > 10 minutes and were assumed to be supporting cells. When advanced carefully forward and out of the presumed supporting cells, the electrode tip entered an extracellular space with zero potential (presumed OoC fluid spaces). When the pipette tip was further advanced by a few microns it encountered cells with resting potentials between - 25 to −45 mV (presumed to be OHCs) that could be held for seconds to several minutes. The electrode tip inevitably encountered the positive potential of the fluid space of the scala media if advanced further through the presumed OHCs.

The electrical resistance between the OoC fluid space immediately adjacent to the OHC basolateral membrane and the scala media was measured by passing 100 Hz sinusoidal current from a calibrated constant current source through a micropipette. The measuring amplifier with the current source was also equipped with a bridge balancing circuit (James Hartley). Most of the micropipette resistance was bridge balanced in the OoC fluid space. This is necessary because the micropipette resistance far exceeds that of the resistance between the scala media and OoC fluid spaces and would dominate any measurements. It was not possible to bridge balance and compensate for the microelectrode resistance completely. Hence, measurements of residual unbalanced microelectrode resistance were made by passing current through the microelectrode while the electrode was still in the OoC fluid spaces. After measurements in the OoC fluid space, the micropipette tip was stepped a few microns through the reticular lamina (RL) into the scala media and the resistance measurement were repeated. The micropipette tip was then stepped back into the OoC fluid space and sinusoidal current was again passed through the tip of the micropipette to check that the micropipette unbalanced resistance remained the same. If the micropipette resistance changed, the data were discarded. The electrical resistance of the current pathway between the OoC fluid space and the scala media was calculated as a difference between the resistance measurements at these two locations. The command voltage for the current injection was software generated and the current source was calibrated by passing it through known value resistors. A wide range of currents from 3×10^−11^ to 5×10^−7^A was used to confirm the linearity of the electrode resistance and the passive resistance between the scala media and the OoC fluid space.

#### Sound system

Sound was delivered via a probe with its tip within 1 mm of the tympanic membrane and coupled to a closed acoustic system comprising two MicroTechGefell GmbH 1-inch MK102 microphones for delivering tones and a Bruel and Kjaer (www.Bksv.co.uk) 3135 0.25-inch microphone for monitoring sound pressure at the tympanum. The sound system was calibrated in situ for frequencies between 1 and 70 kHz by using a laboratory designed and constructed measuring amplifier (James Hartley), and known sound pressure levels (SPLs) were expressed in dB SPL with reference to 2×10^™5^ Pa. White noise for acoustical calibration and tone sequences for auditory stimulation were shaped with raised cosines of 0.5 ms duration were synthesized by a Data Translation 3010 (Data Translation, Marlboro, MA) data acquisition board, attenuated, and used for sound-system calibration and the measurement of electrical cochlear responses.

#### Experimental design and statistical analyses

Heterozygous (CD-1Cx30^A88V/-^) mice were crossed to generate +/+, +/- and -/- genotypes. Male and female mice were studied in approximately equal proportions. No phenotypic differences were observed between males and females. Tests were performed on CD-1Cx30^A88V/A88V^ and CBA/J mice at 3 - 5 weeks of age to standardize our measurements and to reduce animal numbers because of the ease of preparing the cochleae for measurement in this age range. Measurements were made from all CBA/J mice in a litter. Measurements could not effectively be made blind from CD-1 mice with the A88V mutation of connexin30 because homozygous (CD-1Cx30^A88V/A88V^) mice have characteristically rough coats (*14*) and excellent high frequency hearing, whereas smooth coated wild-type littermates have no high frequency hearing and heterozygous littermates are somewhere in between (*15*). Even so, the tested mice were genotyped after each experiment. Experiments were terminated (< 5% of all measurements) if the physiological state of the preparation changed during a measurement in which case data from the sample were excluded. For statistical significance for a given physiological experiment, data were compared from at least 5 CBA/J and 5 CD-1Cx30^A88V/A88V^ mice, obtained from recordings of 2 or 3 complete litters, that were tested before genotypes were determined. Physiological data were plotted as mean ± standard deviation (SD) using Fig P (www.figpsoft.com) or Origin 9.5 (OriginLab Corp. Northampton, MA) software. Statistical tests were performed with GraphPad Prism 8.1 (https://www.graphpad.com/quickcalcs). Statistical comparisons were made using unpaired two-tailed Student’s *t-*tests for unequal variances unless otherwise noted.

### In situ measurements

#### Tissue preparation

The mouse cochlea was freshly isolated and dissected using an ice-cold solution containing in mM: NaCl 135, KCl 5.8, CaCl_2_ 1.3, MgCl_2_ 0.9, NaH_2_PO_4_ 0.7, d-glucose 5.6, HEPES 10, Sodium pyruvate 2, pH 7.5 (adjusted with 1M NaOH), osmolarity ∼ 308 mOsm. DCs were studied in acutely dissected OoC from postnatal day 14 (P14) to P20, where the day of birth is P0. The dissected apical coil of the OoC was transferred to a microscope chamber, immobilized using a nylon mesh fixed to a stainless-steel ring and viewed using an upright microscope (Axioskope, Zeiss, Germany). Cells in the dissected tissue were observed with Nomarski differential interface contrast optics (40× water immersion objectives).

#### Experimental Design and Statistical Analysis

##### Potassium current recordings

All recordings were done at room temperature. Potassium currents were recorded using the same solution as for dissection. Patch pipettes were filled with intracellular solution containing in mM: KCl 131, MgCl_2_ 3, Na_2_ATP 5, Na_2_-phosphocreatine 10, EGTA 1, HEPES 10, Na_2_GTP, pH 7.3 (adjusted with 1M KOH), osmolarity ∼ 293 mOsm. All chemicals were obtained from Sigma Aldrich, UK.

All currents were recorded in a whole-cell voltage-clamp configuration. The patch pipettes were fabricated using borosilicate glass (Sutter Instruments), with 1.5 mm O.D., and heat polished. The pipette resistance in extracellular solution was 2-5 MΩ. Currents were amplified with an Axopatch 700B amplifier (Molecular Devices, Union City, CA) and filtered at a frequency of 2–5 kHz through a low-pass Bessel filter. The data were digitized at 5–500 kHz using an analog-to-digital converter (Digidata 1500; Molecular Devices). The whole-cell current recordings were done using pCLAMP software (version 10, Axon Instruments, Foster City, CA, USA). The sampling frequency was determined by the protocols used. No online leak current subtraction was made, and only recordings with holding currents less than 50 pA were accepted for analyses. The liquid junction potentials were measured (3.5 ± 0.9 mV, n = 149) and corrected online (*68*). The capacitive transients were used to estimate the capacitance of the cell as an indirect measure of cell size. Membrane capacitance was calculated by dividing the area under the transient current in response to a voltage step as described (*69*). The capacitive decay was fitted with a single exponential curve to determine the membrane time constant. Series resistance was estimated from the membrane time constant, given its capacitance. This study included 149 cells with a series resistance (Rs) within a 5 – 15 MΩ range. After 60–90% compensation of the mean residual, uncompensated Rs was 5.1 ± 0.5 MΩ. The seal resistance was typically 2-5 GΩ. The number of cells (n) is given for each data set. Data were analyzed using pClamp10 (Molecular Devices), Origin9.1 (OriginLab Corp. Northampton, MA) and Excel (Microsoft). Time constants (τ_s_) were obtained from fits using Origin software. Time constants were obtained by fitting multiple exponential terms to the activation and decay of the current. The fitting equation was

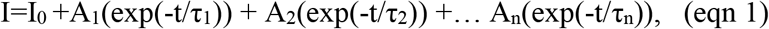

where I_0_ is the initial current magnitude, τ_1_, τ_2_…τ_n_ are the time constants, and A_1_, A_2_…A_n_, are the proportionality constants. The steady-state inactivation curve was generated from normalized currents measured at a test potential following several conditioning pre-pulses. Pooled data were presented as mean ± SD. Statistical comparisons were made using two-tailed Student’s *t-*test. In multiple comparisons tests, significance was adjusted with the Dunn–Sidak correction, where α’ = 1-(1-α)^1/k^, with k the number of comparisons performed between the same groups.

##### Gap junctional conductance recordings

All recordings were done at room temperature. Transjunctional currents were recorded using extracellular solution containing (in mM) NaCl 100, TEA-Cl 20, CsCl 20, BaCl_2_ 1.25, MgCl_2_ 1.48, HEPES 10, pH 7.2 and osmolarity ∼ 300 mOsm. Patch pipettes were filled with the intracellular solution containing in mM: CsCl 140, EGTA 5, MgCl_2_ 2, HEPES 10, pH 7.2, osmolarity ∼ 290 mOsm. Each DC pair was voltage clamped using a dual Axopatch 700B amplifier, (Molecular Devices, Union City, CA) and the response was filtered at a frequency of 2–5 kHz through a low-pass Bessel filter. The data was digitized at 5–500 kHz using an analog-to-digital converter (Digidata 1500; Molecular Devices). The sampling frequency was determined by the protocol used. The liquid junction potentials were measured (4.2 ±0.7 mV, n= 30) and corrected online (*68*). The capacitive transients were used to estimate the capacitance of the cell as an indirect measure of cell size. Membrane capacitance was calculated by dividing the area under the transient current in response to a voltage step as described (*69*). The capacitive decay was fitted with a single exponential curve to determine the membrane time constant. Series resistance was estimated from the membrane time constant, given its capacitance. This study included ∼320 cells with a series resistance (Rs) within a 5– 10 MΩ range. After 60–90% compensation of the mean residual, uncompensated Rs was 4.1±0.6 MΩ. The seal resistance was typically 1–5 GΩ. The transjunctional current recordings were collected using pCLAMP software (version 10, Axon Instruments, Foster City, CA, USA). Data were analyzed using pClamp10 (Molecular Devices), Origin9.1 (OriginLab Corp. Northampton, MA) and Excel (Microsoft).

A classical double voltage clamp technique was used to measure transjunctional conductance (G_j_). Both cells were held at −40 mV. Test voltages ranging from −110 mV to +60 mV were applied to one cell (cell 1) and the other cell (cell 2) was held at a constant voltage of −40 mV. Junctional potential (V_j_) is calculated as

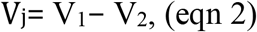

where V_1_ is applied voltage in stepped cell 1 (from −110 to +60 mV), and V_2_ is the holding voltage in non-stepped cell 2 (−40 mV).

Transjunctional currents (I_j_) were measured in a cell 2, as

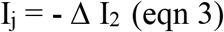

G_j_ was calculated as the slope of the current-voltage relationship by linear regression analysis of the instantaneous current, for the linear I_j_-V_j_ relationship.

For each cell pair, transjunctional conductances (instantaneous, G _j.Inst_ and steady state, G_j.SS_) were calculated as

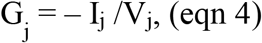

where I_j.Inst_ was measured ∼ 5ms after the begging of the step pulse, and I_j.SS_ was measured at ∼1s at the end of voltage step.

Next, the values for G_j.SS_ and G_j.Inst_ were plotted versus transjunctional voltage V_j_ and the G_j_ – V_j_ curve was fitted with the Boltzmann equation

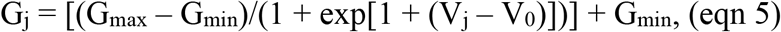

with G_max_ being the maximum conductance (= 1, normalized to instantaneous G_j.Inst_) and G_min_ the minimum conductance. V_0_ is the half inactivation voltage where G_j_ is between G_min_ and G_max_, and A = zq/kT (z = number of equivalent electron charges, q = voltage sensor, k = Boltzmann constant, T = absolute temperature) (*70, 71*).

### Code / software

For physiological recordings, data acquisition and data analysis were performed using a PC with custom programmes written in MATLAB (MathWorks, MA). The programmes are available upon request from authors ANL and IJR. Please note that the programmes were written to communicate with specific hardware (Data Translation 3010 board and custom-made GPIB-controlled attenuators) and will need modifications to be used with different hardware. All proprietary software used in data acquisition and analysis is referred to above in the appropriate section.

## Abbreviations

BM: Basilar membrane
CF: characteristic frequency
Cx26: Connexin26
Cx30: Connexin30
DC: Deiters’ cells
EP: endocochlear potential
ERP: extracellular receptor potential
GJ: Gap junction
MET: Mechano-electrical transducer
OoC: Organ of Corti
OHC: Outer hair cell
RL: Reticular lamina
RP: Receptor potential
TM: Tectorial membrane

## Author contributions

I.J.R., and A.N. L. conceived the research. I.J.R., A.N.L. and S.L. designed the research. S.L. and P.S. performed and analyzed ex vivo physiological measurements. A.N.L. wrote computer programs. V.A.L. and I.J.R. performed and analyzed in vivo physiological recordings. A.N.L. and I.J.R. wrote the paper with significant input from S.L. and P.S.

## Data and materials availability

All data needed to evaluate the conclusions in the paper are present in the paper and/or the Supplementary Materials. Additional data related to this paper may be requested from the authors.

---

**Fig. S1.**
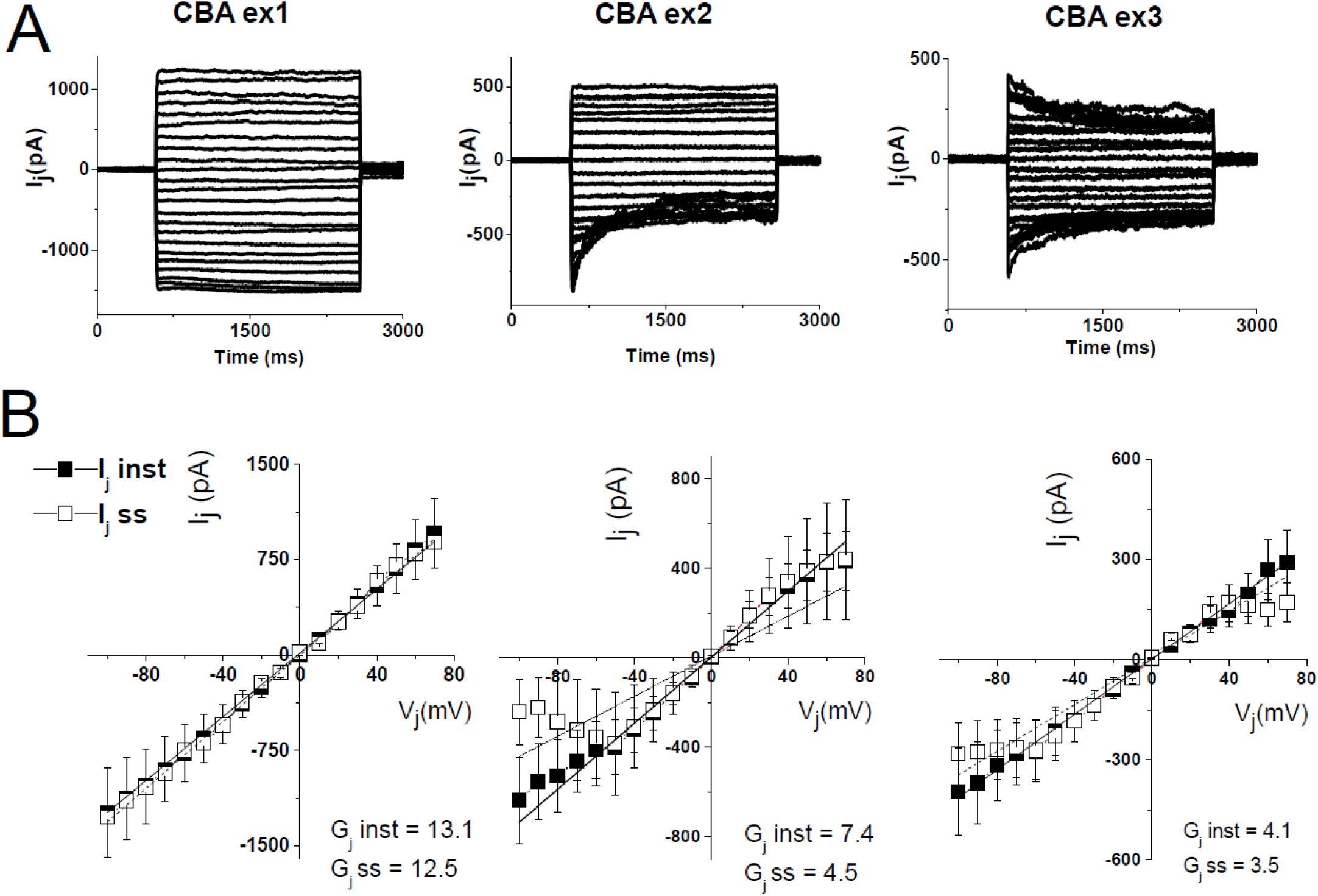
Heterogeneous transjunctional voltage coupling between Deiters cells (DCs) in CBA/J mice. **(A)** Typical examples of transjunctional currents (I_j_) recorded during 2-sec pulse stimulation from pairs of CBA/J DCs. Both cells were initial held at −40 mV and step voltage was applied to one cell and the transjunctional currents were measured in the non-stepped adjacent cell. CBA/J DCs showed a heterogenous transjunctional voltage (V_j_) and time dependence, defined by three different categories of response (HomC, -HetC, ±HetC). **(B)** Summary plots of I_j_ in response V_j_ stimulation obtained from the DC pairs, as classified above. Instantaneous (solid squares) and steady-state (open squares) I_j_ were measured at onset (5 ms) and end (∼2 sec), respectively (showing mean ± SD, n = 6 for each panel). Instantaneous and steady-state gap junctional conductances (G_j_) values correspond to the slope of the linear regression of the represented I_j_-V_j_ relationships.

**Fig. S2.**
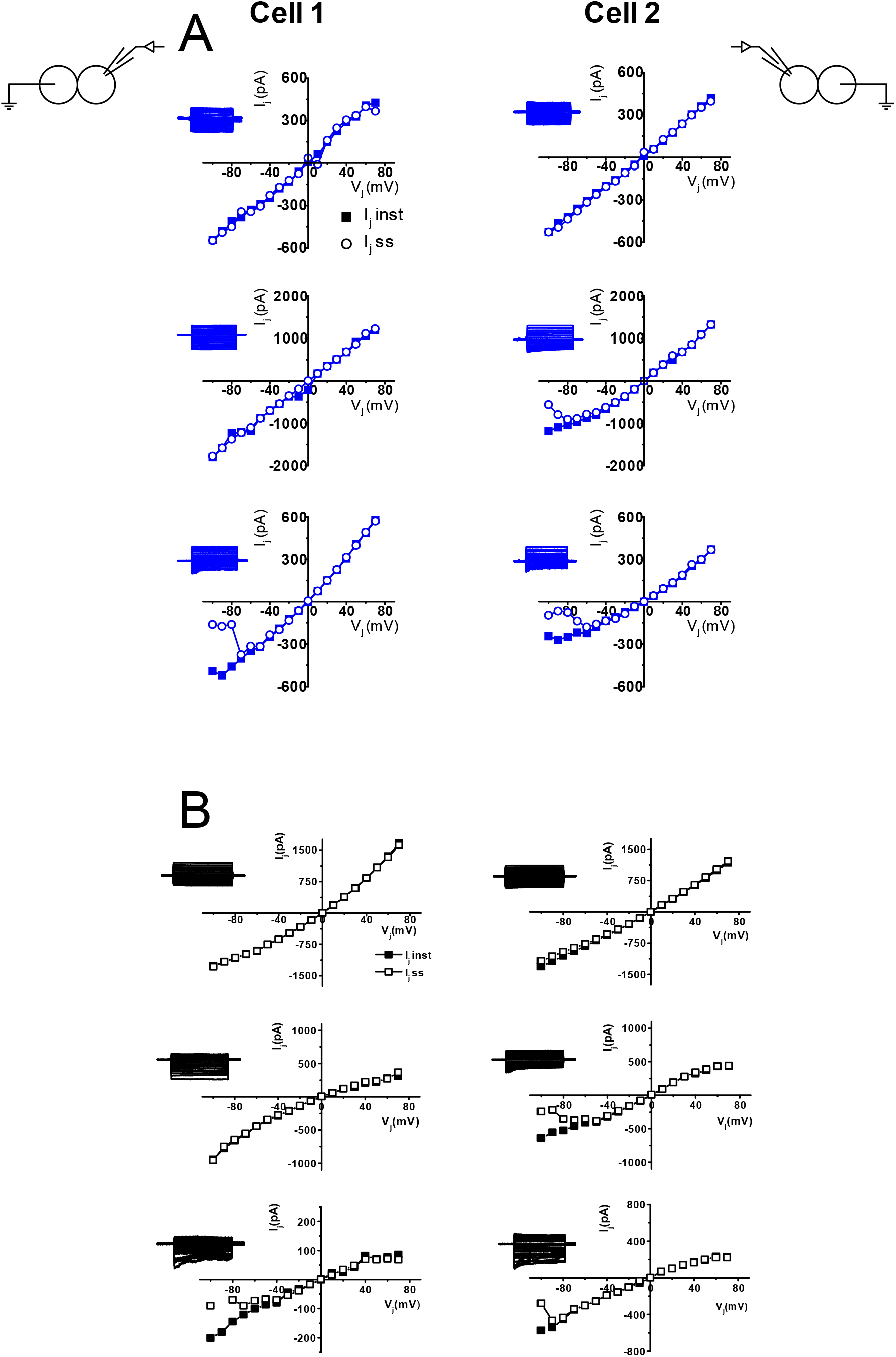
Vj-dependent gating in the same cell pair produces heterogeneous responses in CD-1 (A) and CBA/J (B) Deiters’ cells (DCs). Examples of responses elicited in different coupled DC pairs. (Cell 1) Plots of I_j_ in response to V_j_ stimulation elicited in cell 1 in response to voltage step stimulation of cell 2, and (Cell 2) response of cell 2 to voltage step stimulation in cell 1. Instantaneous (I_j.Inst_, solid squares) and steady-state (I_j.SS_, open circles) were measured at the onset (∼5 ms) and the end of each voltage step (∼2 sec), respectively.

